# GPX4-associated Sedaghatian Type Spondylometaphyseal Dysplasia: A Protein Interactome Perspective

**DOI:** 10.1101/2022.02.17.479371

**Authors:** Kalyani B. Karunakaran, N. Balakrishnan, Madhavi K. Ganapathiraju

## Abstract

**<u>S</u>**pondylo**<u>m</u>**etaphyseal **<u>d</u>**ysplasia, **<u>S</u>**edaghatian type (SMDS) is a rare and lethal skeletal dysplasia inherited in an autosomal recessive manner and caused by mutations in GPX4. In order to expand the functional landscape of this poorly studied disorder and accelerate the discovery of biologically insightful and clinically actionable targets, we constructed SMDS-centric and GPX4-centric protein-protein interaction (PPI) networks, augmented with novel protein interactors predicted by our HiPPIP algorithm. The SMDS-centric networks included those that showed the interconnections of GPX4 with other putative SMDS-associated genes and genes associated with other skeletal dysplasias. The GPX4-centric network showed the interconnections of GPX4 with genes whose perturbation has been known to affect GPX4 expression. We discovered that these networks either contained or were enriched with genes associated with specific SMDS pathophenotypes, tissue-naïve/fetus-specific functional modules and genes showing elevated expression in brain and/or testis similar to GPX4. We identified 7 proteins as novel interactors of GPX4 (APBA3, EGR4, FUT5, GAMT, GTF2F1, MATK and ZNF197) and showed their potential biological relevance to GPX4 or SMDS. Comparative transcriptome analysis of expression profiles associated with chondroplasia and immune-osseous dysplasia versus drug-induced profiles revealed 11 drugs that targeted the neighborhood network of GPX4 and other putative SMDS-associated genes. Additionally, resveratrol, which is currently being tested against a skeletal dysplasia in a clinical trial, was identified as another potential candidate based on the proximity of its targets to GPX4.

## Introduction

**<u>S</u>**pondylometaphyseal **<u>d</u>**ysplasia**<u>s</u>** (SDs) constitute a heterogeneous group of skeletal dysplasias characterized by abnormal development of the spine and the metaphyses of tubular bones, progressive growth and mobility impairment. **<u>S</u>**pondylo**<u>m</u>**etaphyseal **<u>d</u>**ysplasia, **<u>S</u>**edaghatian type (SMDS) is a rare and lethal SD characterized by cupping (or the inward bulging) of the metaphyseal region in long bones (i.e. metaphyseal cupping), flattening of the vertebrae (platyspondyly), abnormal shoulder bone (scapula) morphology, short upper limbs (rhizomelia), abnormal heart rate (arrhythmia), impaired electric conduction from the atrial to the ventricular chamber of the heart (atrioventricular block), cardiorespiratory arrest and several pathologic features in the brain such as the absence/underdevelopment (i.e. agenesis/hypogenesis) of the corpus callosum and underdevelopment of the cerebellum (cerebellar hypoplasia).^1–8^ Respiratory distress is the primary cause of death for infants born with this congenital condition.^1,7,8^ Four patients living with this debilitating disorder exhibit delayed cognitive development, severe hypotonia characterized by a lack of neck muscle control and an inability to sit or walk unsupported, and intractable epileptic seizures.^9^ SMDS shows an autosomal recessive pattern of inheritance, and has been attributed to at least 3 mutations in the gene GPX4 (located in the chromosomal region 19p13.3), which codes for the protein *glutathione peroxidase*, namely, c.381C>A (p.Tyr90Ter/Y90*) (single nucleotide variant in exon 3 that introduces a translation stop before the indicated amino acid position in the protein), c.587+5G>A (single nucleotide variant in intron 4 causing splicing out of a part of exon 4, indicated here with reference to the nearest coding sequence) and c.588-8_588-4del (5 bp deletion in intron 4 causing skipping of exon 5, indicated here with reference to the nearest coding sequence).^7,8,10^ GPX4 is a multifunctional (‘moonlighting’) protein that not only catalyzes the reduction of lipoproteins and membrane phospholipids to prevent oxidative damage resulting from lipid peroxidation,^11^ but also serves as a structural protein in the midpiece of mature sperm.^12^ GPX4 activity and synthesis are influenced by Selenium levels, as its catalytic site contains a selenocysteine residue.^13^ Therapeutic options for SMDS are dismally limited to controlling oxidative damage using vitamin E, n-acetyl-cysteine and coenzyme Q10 supplements. Except for some early findings on the clinical presentation of SMDS^1–6^ and the characterization of the GPX4 mutations,^7,8^ the etiology of this severe lethal dysplasia remains largely unexplored. Hence, novel approaches are needed to elucidate the broad themes underlying this disorder.

Protein-protein interactions (PPIs) drive the cellular machinery by facilitating a variety of biological processes including signal transduction, formation of cellular structures and enzymatic complexes. Disease-associated variants are enriched in protein cores and protein interaction interfaces.^14^ Variants localized to the protein core may disrupt its tertiary structure and abolish all chances of the protein interacting with any of its interaction partners (node removal in the interactome).^14^ Variants localized to interaction interfaces may perturb specific interactions (edge perturbation in the interactome).^14^ Over two-thirds of disease-associated variants alter binding affinity or even establish novel interactions.^15^ Hence, genetic mutations may perturb proteins and this effect may spread in the PPI network (or the ‘interactome’) affecting other proteins, posing deeper implications for disease development,^16^ for example multiple pathophenotypes in a single disease despite the disease being associated with a single genotype.^17^ Several studies, led by our own group and others, have successfully traced shared genetics and symptomatology among different diseases back to this complex network of PPIs.^18–21^ Therefore, for functional interpretation of genetic variants associated with complex disorders, it is imperative that we place variants in the complex web of PPIs. Mapping disease-associated variants onto PPI networks will pull in more disease-associated genes into the network, offering an opportunity to analyze communities of proteins involved in mechanisms relevant to disease etiology.

In this study, we adopted an interactome-based approach to construct an integrated functional landscape for GPX4-associated SMDS in relation with other SDs, phenotypically similar dysplasias, genes other than GPX4 speculated to be associated with SMDS and genes whose perturbation affects GPX4 expression. We identified functional modules and pathophenotypes enriched in this landscape, and used human transcriptomic data to identify groups of genes clustering with GPX4, which showed elevated expression in specific tissues. Finally, we adopted two approaches to propose a few repurposable drugs, namely, comparative analysis of drug-induced and disease-associated transcriptomes, and examining the network proximity of GPX4 to the protein targets of drugs that are being used or being tested against other skeletal dysplasias.

## Results

We assembled the known PPIs of GPX4 from HPRD^22^ (Human Protein Reference Database) and BioGRID^23^ (Biological General Repository for Interaction Datasets), and predicted its novel PPIs by applying the HiPPIP algorithm described in our earlier work.^20^ HiPPIP computes features of protein pairs such as cellular localization, molecular function, biological process membership, genomic location of the gene, and gene expression in microarray experiments, and classifies the pairwise features as *interacting* or *non-interacting* based on a random forest model.^20^ HiPPIP was shown to have high precision in its original evaluation^20^ and to be superior to state-of-the-art algorithms in a recent evaluation.^24^ Eighteen novel PPIs predicted by HiPPIP were experimentally tested and all eighteen were found validated to be true PPIs by collaborators as summarized in our recent work.^25^ GPX4 has two known interactions, with MAPK13 (*mitogen-activated protein kinase 13*) and PRDX6 (*peroxiredoxin 6*). Additionally, we predicted seven novel interactions with APBA3 (*amyloid beta precursor protein binding family A member 3*), EGR4 (*early growth response 4*), FUT5 (*fucosyltransferase 5*), GAMT (*guanidinoacetate N-methyltransferase*), GTF2F1 (*general transcription factor IIF subunit 1*), MATK (*megakaryocyte-associated tyrosine kinase*) and ZNF197 (*zinc finger protein 197*). **Table 1** lists the evidence supporting the biological relevance of these novel interactors to GPX4 or SMDS, identified from our analyses.

**Table 1:**
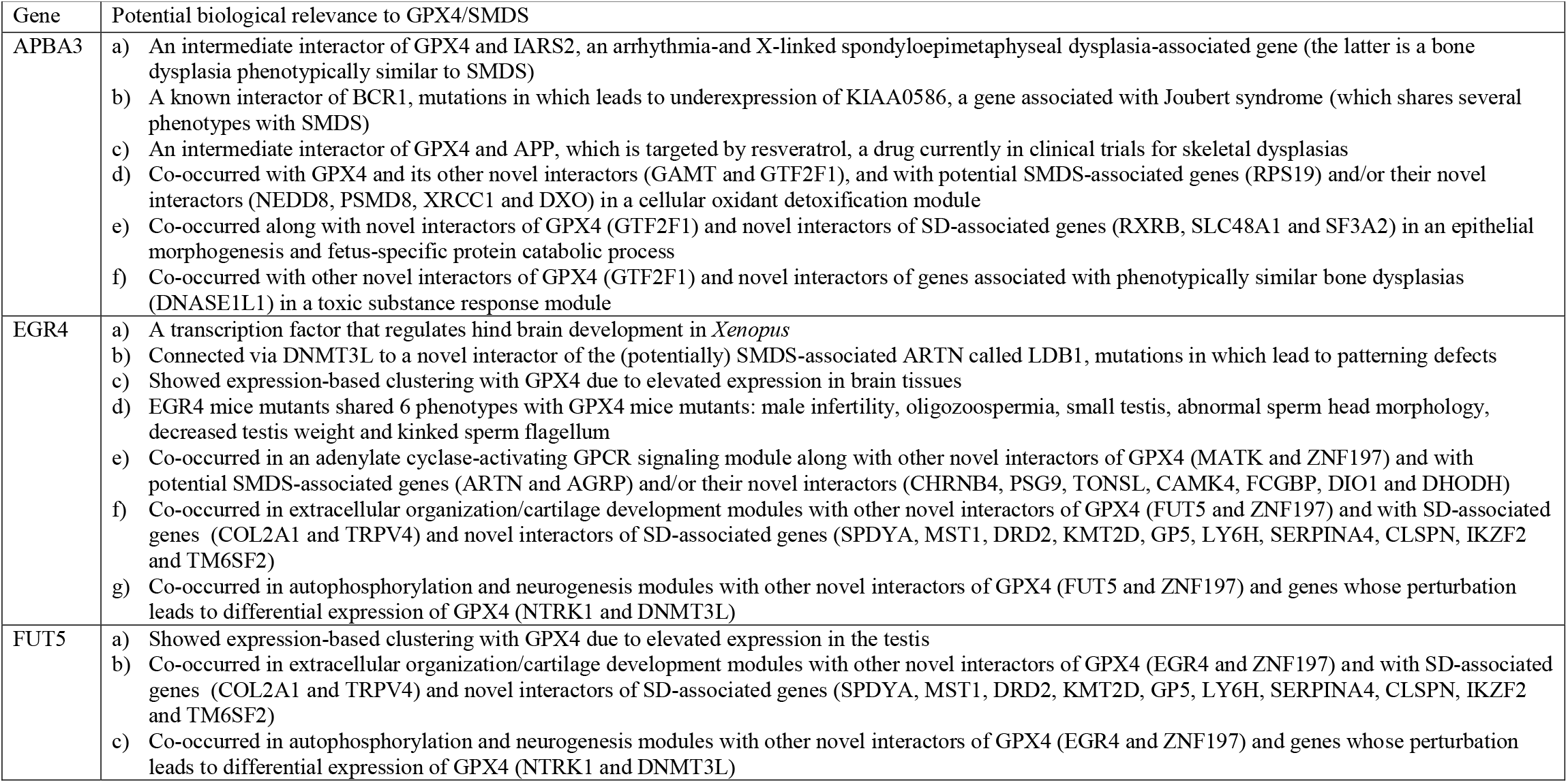
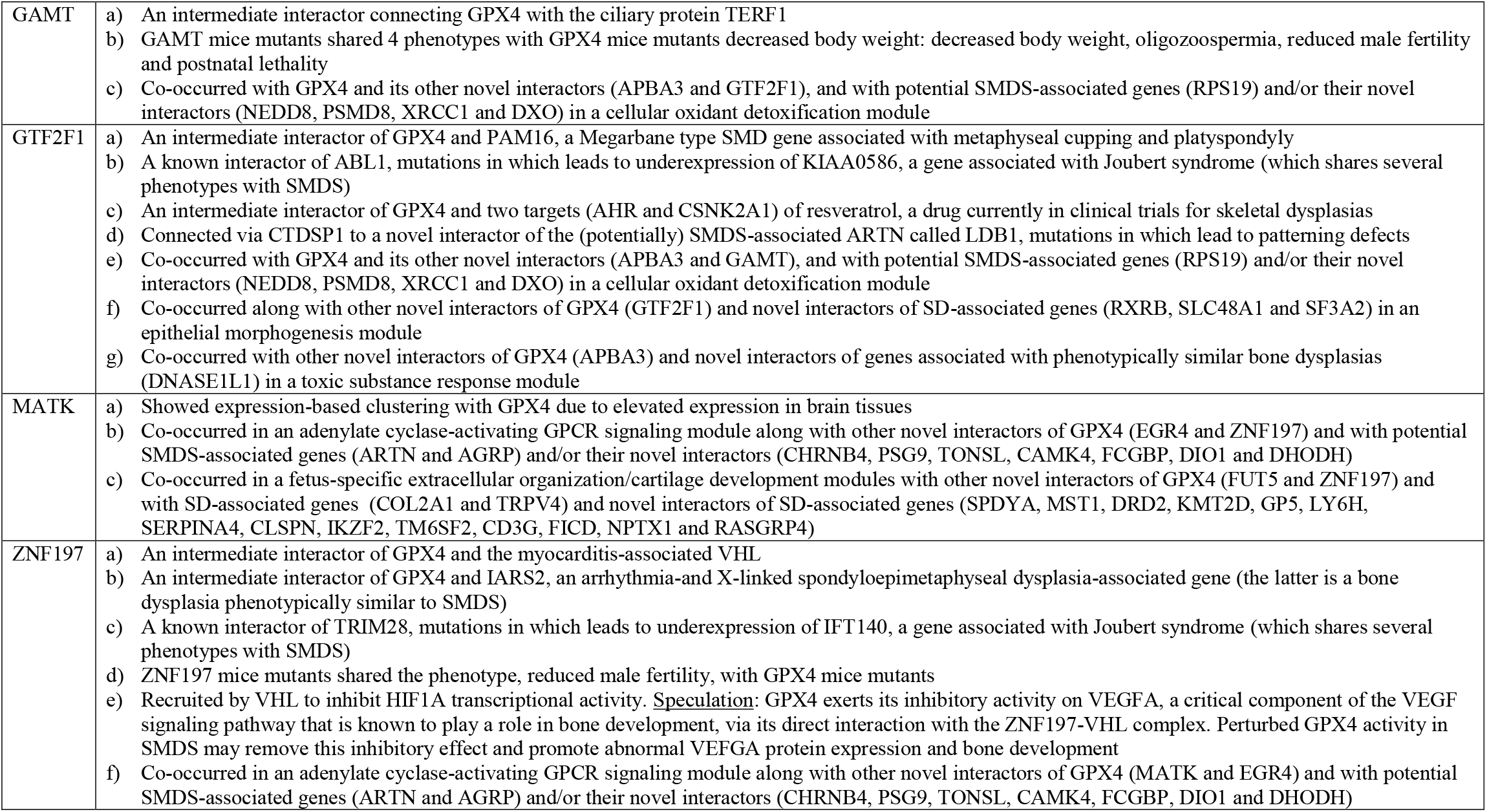
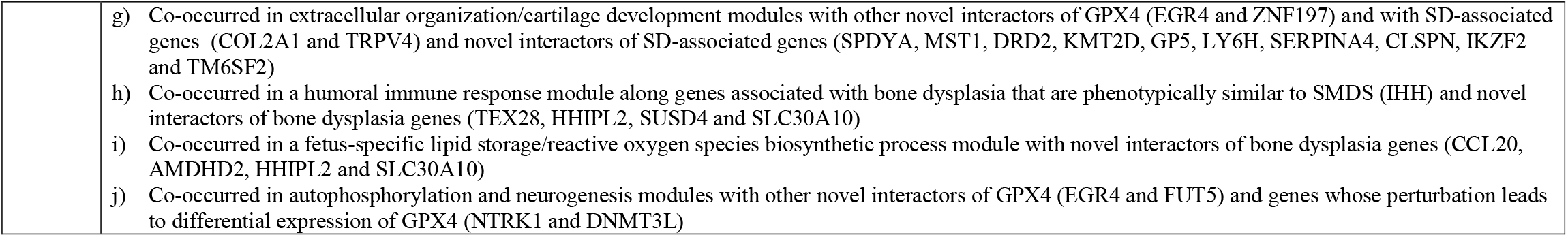
Biological relevance of novel interactors of GPX4. The table lists different pieces of evidence gathered from our study that support the biological validity of the computationally predicted novel interactors of GPX4 to SMDS or GPX4 itself. Note that in the table, “phenotypic similarity” refers to “phenotypic similarity with SMDS” (unless specified otherwise). SMDS: SpondyloMetaphyseal Dysplasia, Sedaghatian type, SD: Spondylometaphyseal Dysplasia.

Protein interactome analysis is a useful tool for elucidating biologically relevant relationships existing at a higher level among genes, which may not be apparent by examining individual genes. Therefore, for further mechanistic characterization of GPX4, we inspected its interconnections in the human interactome with (a) other genes speculated to be associated with SMDS in the DisGeNET^26^ database, (b) genes associated with other types of SDs, (c) genes associated with disorders that shared phenotypic similarity with SMDS and (d) genes whose perturbation was known to cause significant overexpression or underexpression of GPX4. Following this, we used the HumanBase toolkit^27^ to identify functional modules in each of these GPX4-associated interactomes. HumanBase employs shared k-nearest-neighbors and the Louvain community-finding algorithm to cluster the genes sharing the same network neighborhoods and similar GO biological processes into functional modules. Additionally, we compiled the genes involved in 12 major pathophenotypes associated with SMDS from the MONARCH^28^ database and computed the statistical significance of the enrichment of these genes in each of the interactomes, against a background of 15451 genes with phenotype associations. The selected phenotypes were atrial septal defect (290 genes), atrioventricular block (54 genes), cardiorespiratory arrest (10 genes), cerebellar hypoplasia (219 genes), myocarditis (7 genes), agenesis of corpus callosum (240 genes), arrhythmia (355 genes), pachygyria (133 genes; cerebral cortex malformation characterized by a fewer number of abnormally wide gyri), metaphyseal cupping (18 genes), platyspondyly (109 genes), abnormal scapula morphology (4 genes) and abnormality of the ribs (85 genes).

### Interactome of GPX4 and other genes potentially associated with SMDS

The human sperm contains 648 short exon-sized sequences called sperm RNA elements corresponding to a range of unique coding and non-coding transcripts, the presence of which may increase the likelihood of live birth through natural conception in idiopathic infertile couples.^29^ A sperm RNA element that was mapped to GPX4 contained an SMDS-associated variant.^29^ The DisGeNET^26^ database lists 6 additional SMDS-associated genes extracted from this work by the BeFree^30^ text mining system (i.e. other than GPX4 which had a gene-disease association or GDA score of 0.72), namely, AGRP (agouti related neuropeptide), ARNTL (aryl hydrocarbon receptor nuclear translocator like), ARTN (artemin), LOH19CR1 (loss of heterozygosity, 19, chromosomal region 1), PSD4 (pleckstrin and Sec7 domain containing 4) and RPS19 (ribosomal protein S19) (GDA score = 0.01). BeFree employs a kernel-based approach based on morphosyntactic and dependency information to identify gene-disease associations.^30^ We extracted the PPI network that connects these 6 genes (candidates) to GPX4 (target) through shortest paths as well as their own interactors (**Fig. 1**), and found that GPX4 connects to AGRP, ARNTL, ARTN, PSD4 and RPS19 through 22 intermediate interactors including 10 novel interactors (i.e. those revealed by computationally predicted PPIs in this work). These novel interactors include 6 novel interactors of GPX4 (APBA3, EGR4, GAMT, GTF2F1, MATK and ZNF197), a direct interactor of ARTN (LDB1), a direct interactor of AGRP (UPP1) and 2 novel intermediate interactors connecting GPX4 with AGRP (VHL and GHRL).

**Figure 1:**
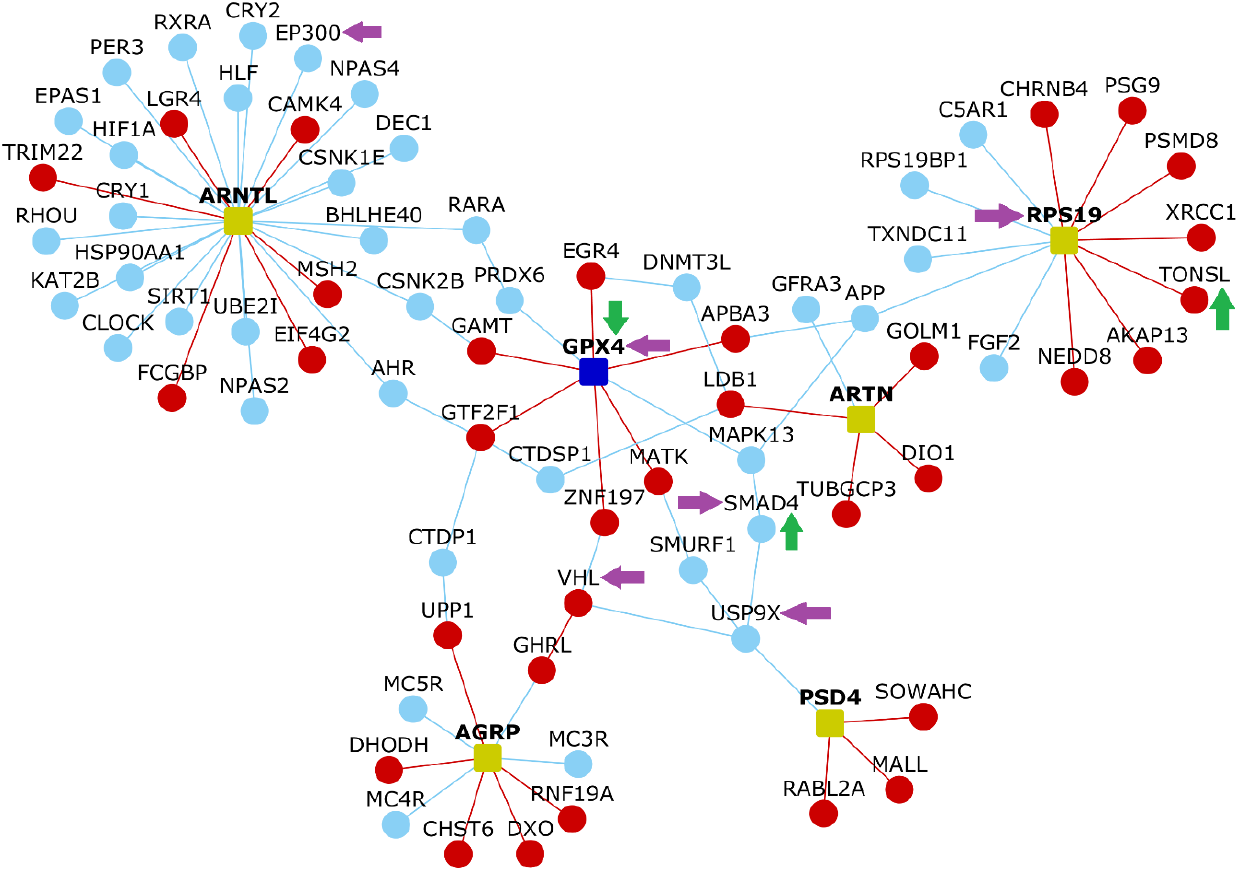
Interactions of GPX4 with other genes speculated to be associated with Sedaghatian type spondylometaphyseal dysplasia. This network diagram shows the shortest paths connecting the genes from the DisGeNET database (dull green colored nodes) identified through text mining to GPX4. Nodes depict proteins and edges depict PPIs. Red and light blue colored nodes denote novel and known interactors respectively. Red and light blue colored edges denote novel and known PPIs respectively. Purple colored arrows indicate genes associated with cardiac defects, whereas green colored arrows indicate those associated with skeletal defects.

This interactome was enriched in genes associated with myocarditis (*P*-value = 5.33E-04, odds ratio = 55.8, genes: VHL and GPX4), atrial septal defect (*P*-value = 0.016, odds ratio = 3.37, genes: USP9X, SMAD4, GPX4, RPS19 and EP300) and platyspondyly (*P*-value = 0.018, odds ratio = 5.38, genes: GPX4, SMAD4 and TONSL). Myocarditis-associated VHL was connected to GPX4 via ZNF197. Platypondyly-associated TONSL was predicted to be a novel interactor of RPS19.

Using the HumanBase toolkit,^27^ we isolated the functional modules enriched in the interactome containing the intermediate interactors connecting GPX4 to the additional 6 potential SMDS-associated genes, as well as the direct known and novel interactors of these 6 genes. We extracted tissue-specific functional modules containing genes specific to the human fetal tissue and tissue-naive (or ‘global’) modules containing genes playing identical roles across the tissues. Fetus-specific modules were examined based on the assumption that modules putatively active in the fetal tissue may be relevant to the congenital and neonatal underpinning of SMDS. A *cellular oxidant detoxification* module was identified in both global (M4: *Q*-value = 2.15E-03) and fetus-specific (M3: *Q*-value = 2.77E-03) contexts. GPX4 and 3 of its novel interactors (APBA3, GAMT and GTF2F1), RPS19 and 3 of its novel interactors (NEDD8, PSMD8 and XRCC1) and a novel interactor of AGRP called DXO were detected in the global functional module. The fetus-specific module did not contain DXO, but instead contained a novel interactor of ARTN called LDB1. The *adenylate cyclase-activating G-protein coupled receptor signaling pathway* was detected both as a global (M3: *Q*-value = 4.25E-04) and fetus-specific (M2: *Q*-value = 3.4E-04) module. 3 novel interactors of GPX4 (EGR4, MATK and ZNF197), 3 novel interactors of RPS19 (CHRNB4, PSG9 and TONSL), 2 novel interactors of ARNTL (CAMK4 and FCGBP), ARTN and its novel interactor DIO1, and AGRP and its novel interactor DHODH were detected in the global module. The fetus-specific module did not contain AGRP and DHODH, but instead contained VHL, a novel intermediate interactor between GPX4 and AGRP. *Cell growth* (M1: *Q*-value = 2.38E-04) and *response to redox state* (M2: *Q*-value = 2.38E-04) were detected only as global modules. The *cell growth module* contained 2 novel interactors of ARNTL (EIF4G2 and MSH2) and a novel interactor of ARTN (TUBGCP3). The *redox state response* module contained a novel interactor of AGRP called UPP1. *N-terminal peptidyl—lysine acetylation* (M1: *Q*-value = 1.21E-04)/*histone acetylation* (M1: *Q*-value = 2.84E-04) and *regulation of DNA metabolic process* (M4: *Q*-value = 7.84E-04) were detected only as tissue-specific modules. A novel interactor of ARNTL (TRIM22) and a novel interactor of AGRP (RNF19A) belonged to the *acetylation* module, whereas MSH2 and TUBGCP3 that were earlier detected in the *cell growth* module, seemed to also belong to the *DNA metabolic process* module.

### Interactome of GPX4 and genes associated with other spondylometaphyseal dysplasias

We curated genes associated with other types of SDs from a comprehensive review on genetic skeletal disorders^31^ in order to map their connections to GPX4 in the human interactome. Specifically, this included a set of 9 genes associated with Kozlowski type SMD (TRPV4), spondyloenchondrodysplasia (ACP5), odontochondrodysplasia (TRIP11), Sutcliffe/corner fractures type SMD, (FN1 and COL2A1), SMD with severe genu valgam/Schmidt type SMD (COL2A1), SMD with cone-rod dystrophy (PCYT1A), SMD with retinal degeneration/axial SMD (CFAP410), dysspondyloenchondromatosis (COL2A1), achondrogenesis type 1A (TRIP11), schneckenbecken dysplasia (SLC35D1 and INPPL1) and opsismodysplasia (INPPL1). We extracted the shortest paths connecting these 9 genes (candidates) to GPX4 (target). GPX4 shared 93 intermediate interactors with these 9 genes, including 26 novel interactors (**Fig. 2**). These novel interactors included 4 direct interactors of GPX4 (APBA3, GTF2F1, MATK and ZNF197), 2 of CFAP410/C21orf2 (COL6A1 and COL6A2), 3 of TRIP11 (ASB2, EPS8 and SERPINA4), 2 of ACP5 (EPOR and SF3A2), 2 of SLC35D1 (MACF1 and VCAM1), 2 of PCYT1A (PAK2 and RUBCN), 1 each of FN1 (CREB1), TRPV4 (RXRB) and COL2A1 (DCN) and 8 intermediate interactors (RPS6KA2, PELI2, THRA, STAT5A, SKIC, TNFSF10, PLAT and PPP2CB). GPX4 was more closely connected with the genes associated with Kozlowski type SMD (TRPV4), odontochondrodysplasia, achondrogenesis type 1A (TRIP11), Sutcliffe type SMD (FN1), SMD with cone-rod dystrophy (PCYT1A), schneckenbecken dysplasia and opsismodysplasia (INPPL1).

**Figure 2:**
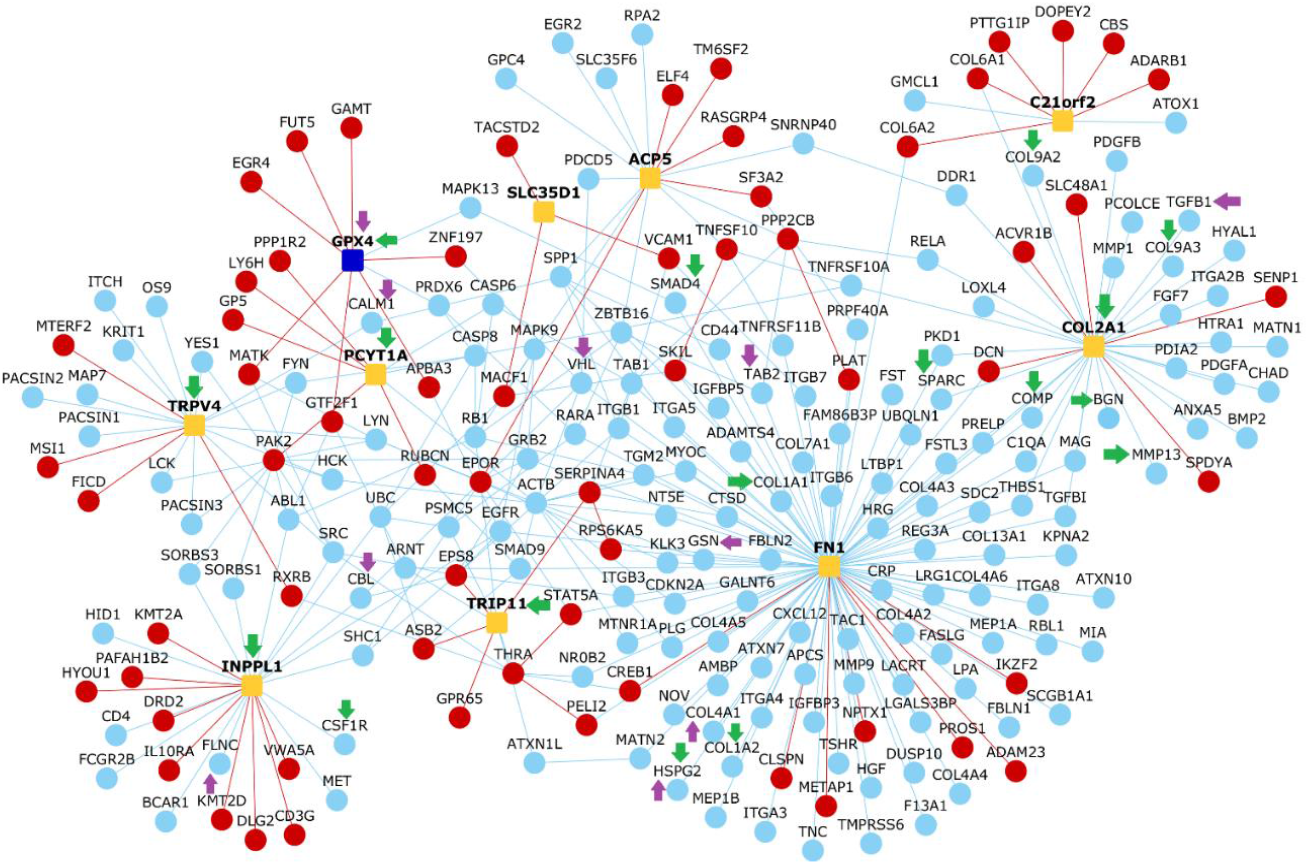
Interactions of GPX4 with genes associated with other spondylometaphyseal dysplasias. This network diagram shows the shortest paths connecting GPX4 with the genes associated with Kozlowski type SMD, spondyloenchondrodysplasia, odontochondrodysplasia, Sutcliffe/corner fractures type SMD, SMD with severe genu valgam/Schmidt type SMD, SMD with cone-rod dystrophy, SMD with retinal degeneration/axial SMD, dysspondyloenchondromatosis, achondrogenesis type 1A, schneckenbecken dysplasia and opsismodysplasia (orange colored nodes). Nodes depict proteins and edges depict PPIs. Red and light blue colored nodes denote novel and known interactors respectively. Red and light blue colored edges denote novel and known PPIs respectively. Purple and green arrows indicate genes associated with cardiac and skeletal defects respectively.

This interactome was enriched in genes associated with arrhythmia (*P*-value = 0.041, odds ratio = 1.88, genes: TAB2, COL4A1, HSPG2, FLNC, CALM1, GPX4, CBL, VHL, GSN and TGFB1), cardiorespiratory arrest (*P*-value = 9.25E-03, odds ratio = 13.38, genes: COL2A1 and GPX4), myocarditis (*P*-value = 4.64E-03, odds ratio = 19.11, genes: GPX4 and VHL), metaphyseal cupping (*P*-value = 5.9E-03, odds ratio = 3.68, genes: GPX4, PCYT1A, INPPL1, TRIP11, MMP13 and COL2A1), platyspondyly (*P*-value = 4.98E-13, odds ratio = 10.43, genes: GPX4, PCYT1A, INPPL1, TRIP11, MMP13, COL2A1, HSPG2, SPARC, BGN, COMP, COL1A1, COL9A2, COL9A3, CSF1R, TRPV4, COL1A2 and SMAD4) and abnormal ribs (*P*-value = 8.9E-03, odds ratio = 3.93, genes: GPX4, PCYT1A, HSPG2, TRPV4 and SMAD4).

We examined the functional modules enriched in this interactome, which contained the intermediate interactors connecting GPX4 to the genes associated with other SDs, and the known and novel interactors of these genes. Three pairs of closely related modules were detected in global and fetal contexts: *blood vessel development* (M1: *Q*-value = 9.6E-06)/*positive regulation of cell migration* (M1: *Q*-value = 2.77E-06), *extracellular organization* (M3: *Q*-value = 3.53E-04)/*cartilage development* (M3: *Q*-value = 2.72E-03) and *regulation of cysteine-type endopeptidase activity involved in apoptotic process* (M6: *Q*-value = 6.68E-03) *execution phase of apoptosis* (M4: *Q*-value = 6.75E-04). Three unique global modules were identified: *leukocyte tethering or rolling* (M2: *Q*-value = 3.7E-05), *response to bacterium* (M4: *Q*-value = 6.4E-03) and *morphogenesis of an epithelium* (M5: *Q*-value = 6.57E-03). Four unique fetus-specific modules were identified: *cellular component maintenance* (M2: *Q*-value = 3.39E-04), *actin filament organization* (M5: *Q*-value = 1.36E-03), *negative regulation of intrinsic apoptotic signaling pathway* (M6: *Q*-value = 2.51E-03) and *regulation of protein catabolic process* (M7: *Q*-value = 1.05E-02). 3 novel interactors of GPX4 (EGR4, FUT5 and ZNF197) co-occurred with 10 novel interactors of genes associated with other SD genes (SPDYA, MST1, DRD2, KMT2D, GP5, LY6H, SERPINA4, CLSPN, IKZF2 and TM6SF2) and 2 SD genes themselves (COL2A1 and TRPV4) in the *extracellular organization*/*cartilage development* modules in both global and fetus-specific contexts. The global module additionally contained the novel interactors THRA and ADAM23, and the fetus-specific module contained MATK (a novel interactor of GPX4), CD3G, FICD, NPTX1 and RASGRP4. 2 novel interactors of GPX4 (APBA3 and GTF2F1) co-occurred with 3 novel interactors of SD-associated genes (RXRB, SLC48A1 and SF3A2) in the *epithelial morphogenesis* (global) module. APBA3, a novel interactor of GPX4, co-occurred with 2 novel interactors of SD-associated genes (RXRB and SF3A2) in the *protein catabolic process* (fetus-specific) module.

### Interactome of GPX4 and genes associated with disorders sharing phenotypic similarity with SMDS

We employed Phenogrid from the MONARCH toolkit^28^ to identify the diseases that were most phenotypically similar to SMDS. The phenogrid algorithm identifies the common phenotypes between two given diseases. It then assesses the information content in each of these phenotypes (gene and disease associations) to assign a specific strength to the similarity observed between the diseases. Seven bone dysplasias appeared to be sharing more than 70% phenotypic similarity with SMDS, namely, X-linked spondyloepimetaphyseal dysplasia (77%), acrocapitofemoral dysplasia (75%), A4 type SMD (75%), Dyggve-Melchior-Clausen disease (74%), autosomal recessive Megarbane type SMD (73%), metaphyseal acroscyphodysplasia (72%) and X-linked Dyggve-Melchior-Clausen disease (71%). We checked whether genetic associations were available for these 7 dysplasias in DisGeNET.^26^ 4 of them appeared to have been either causally linked or correlated with a few genes, namely, X-linked spondyloepimetaphyseal dysplasia with BGN and IARS2, acrocapitofemoral dysplasia with IHH, Dyggve-Melchior-Clausen disease with DYM and autosomal recessive Megarbane type SMD with PAM16. Next, we extracted the shortest paths connecting these 5 genes (candidates) to GPX4 (target). GPX4 was found to be connected to these 5 genes through 13 intermediate interactors including 5 novel interactors, namely, 3 novel interactors of GPX4 (APBA3, GTF2F1 and ZNF197) and 2 novel interactors of DYM (EPG5 and MAPK4) (**Fig. 3**).

**Figure 3:**
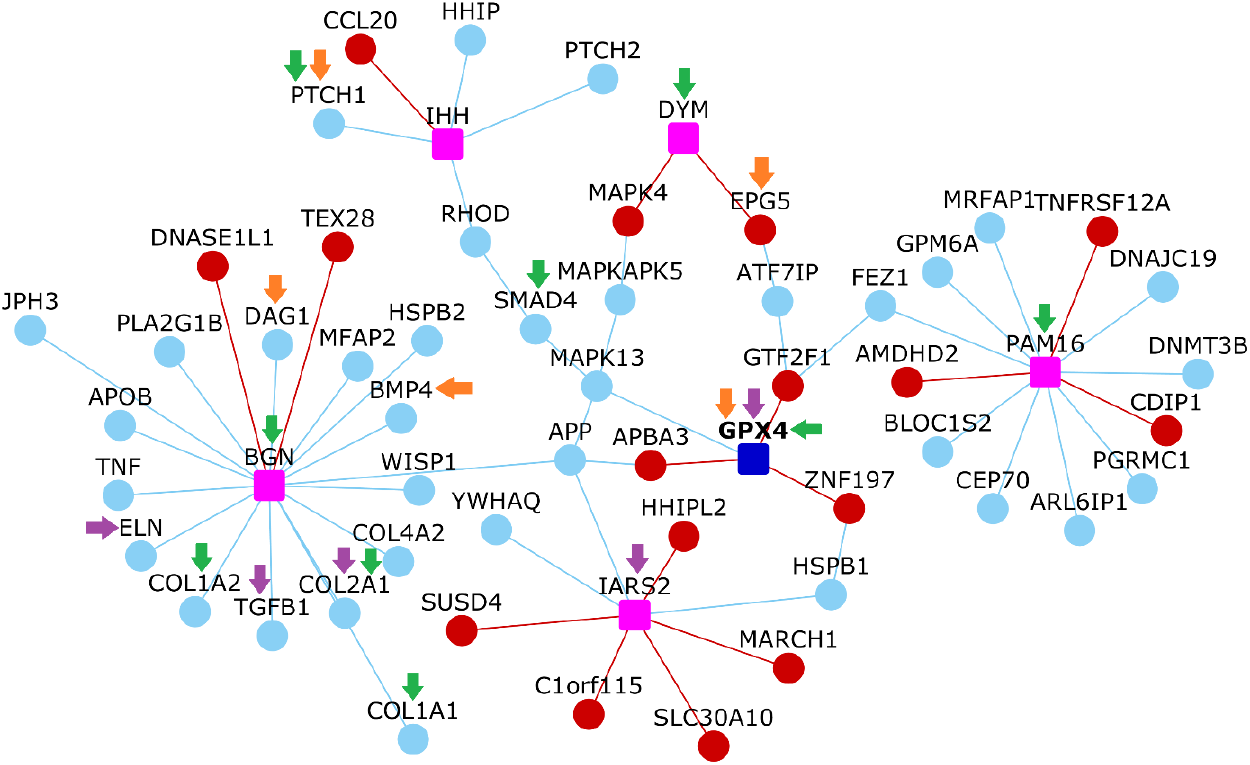
Interactions of GPX4 with genes associated with bone dysplasias sharing phenotypic similarity with Sedaghatian type spondylometaphyseal dysplasia. This network diagram shows the shortest paths connecting GPX4 with the genes associated with X-linked spondyloepimetaphyseal dysplasia, acrocapitofemoral dysplasia, Dyggve-Melchior-Clausen disease and autosomal recessive Megarbane type SMD (pink colored nodes). Nodes depict proteins and edges depict PPIs. Red and light blue colored nodes denote novel and known interactors respectively. Red and light blue colored edges denote novel and known PPIs respectively. Purple, green and orange arrows indicate genes associated with cardiac, skeletal and brain defects respectively.

This interactome was enriched in genes associated with arrhythmia (*P*-value = 0.042, odds ratio = 3.05, genes: TGFB1, IARS2, GPX4 and ELN), cardiorespiratory arrest (*P*-value = 5.9E-04, odds ratio = 54.2, genes: COL2A1 and GPX4), cerebellar hypoplasia (*P*-value = 8.63E-03, odds ratio = 4.95, genes: BMP4, EPG5, GPX4 and DAG1), agenesis of corpus callosum (*P*-value = 1.88E-03, odds ratio = 5.65, genes: BMP4, EPG5, GPX4, PTCH1 and DAG1), metaphyseal cupping (*P*-value = 3.73E-05, odds ratio = 45.18, genes: GPX4, COL2A1 and PAM16), platyspondyly (*P*-value = 5.9E-09, odds ratio = 19.9, genes: GPX4, COL2A1, PAM16, BGN, DYM, SMAD4, COL1A1 and COL1A2) and abnormal ribs (*P*-value = 3.8E-03, odds ratio = 9.57, genes: GPX4, SMAD4 and PTCH1). Arrhythmia-associated IARS2 is associated with X-linked spondyloepimetaphyseal dysplasia and separated from GPX4 only by 3 edges and 5 intermediate interactors (including 2 novel interactors of GPX4, namely, APBA3 and ZNF197). EPG5, the gene associated with cerebellar hypoplasia and corpus callosum agenesis, is a novel interactor of DYM, a Dyggve-Melchior-Clausen disease gene, and is an intermediate interactor connecting DYM to GPX4. PAM16, associated with metaphyseal cupping and platyspondyly, is a Megarbane type SMD gene, and is separated from GPX4 only by 3 edges and 2 intermediate interactors (including the novel interactor of GPX4, GTF2F1). Both BGN associated with X-linked spondyloepimetaphyseal dysplasia and DYM associated with Dyggve-Melchior-Clausen disease were separated from GPX4 by 3 edges, and appeared to be linked to platyspondyly.

Five modules were detected in both global and fetus-specific contexts from the interactome: *skeletal system development* (global M1: *Q*-value = 2.76E-06)/*embryo development/gastrulation* (fetus-specific M1: *Q*-value = 2.28E-04), *lipid storage* (global M2: *Q*-value = 1.69E-04)/*reactive oxygen species biosynthetic process* (fetus-specific M2: *Q*-value = 9.17E-04), *humoral response* (global M3: *Q*-value = 2E-03 and fetus-specific M3: *Q*-value = 1.28E-03), *cellular response to toxic substance* (global M4: *Q*-value = 2.59E-03 and fetus-specific M4: *Q*-value = 3.76E-03) and *membrane organization* (global M5: *Q*-value = 8.47E-03 and fetus-specific M6: *Q*-value = 6.29E-03). We noted another *embryo development* module that was fetus-specific and enriched for *neurogenesis* (M5: *Q*-value = 0.01). GPX4 and 2 of its novel interactors (APBA3 and GTF2F1) co-occurred with a novel interactor of a gene associated with a bone dysplasia (DNASE1L1) in the *toxic substance response* module in both global and fetus-specific contexts. 1 novel interactor (AMDHD2) and 2 novel interactors (CDIP1 and C1orf115) uniquely occurred in this module in global and fetus-specific contexts respectively. ZNF197, a novel interactor of GPX4, co-occurred with 4 novel interactors of bone dysplasia genes (TEX28, HHIPL2, SUSD4 and SLC30A10) and a bone dysplasia gene itself (IHH) in the global *humoral immune response* module. ZNF197 also co-occurred with 4 novel interactors of bone dysplasia genes (CCL20, AMDHD2, HHIPL2 and SLC30A10) in the fetus-specific *lipid storage/reactive oxygen species biosynthetic process* module.

### Interactome of GPX4 and genes whose perturbation affects GPX4 expression

Using the Knockdown Atlas from the BaseSpace Correlation Engine software suite,^32^ we compiled a list of 136 genes whose perturbation, in the form of gene knockout/knockdown/mutation/overexpression, significantly affects GPX4 expression. We extracted the shortest paths connecting these 136 genes (candidates) to GPX4 (target). Out of the 136 genes, 4 were closely connected to GPX4, namely, AHR, DNMT3L, JUN and NTRK1, through 33 intermediate interactors including 9 novel interactors (**Fig. 4**). These novel interactors included all the 7 novel interactors of GPX4 (APBA3, EGR4, FUT5, GAMT, GTF2F1, MATK and ZNF197) and 2 intermediate interactors (MAPK13 and SUMO1). Mutations in JUN lead to GPX4 underexpression, whereas DNMT3L knockdown, AHR knockout and NTRK1 overexpression lead to GPX4 overexpression. This interactome was enriched in genes associated with cardiorespiratory arrest (*P*-value = 3.2E-03, odds ratio = 23.06, genes: GPX4 and KIT) and myocarditis (*P*-value = 1.52E-03, odds ratio = 32.9, genes: GPX4 and VHL).

**Figure 4:**
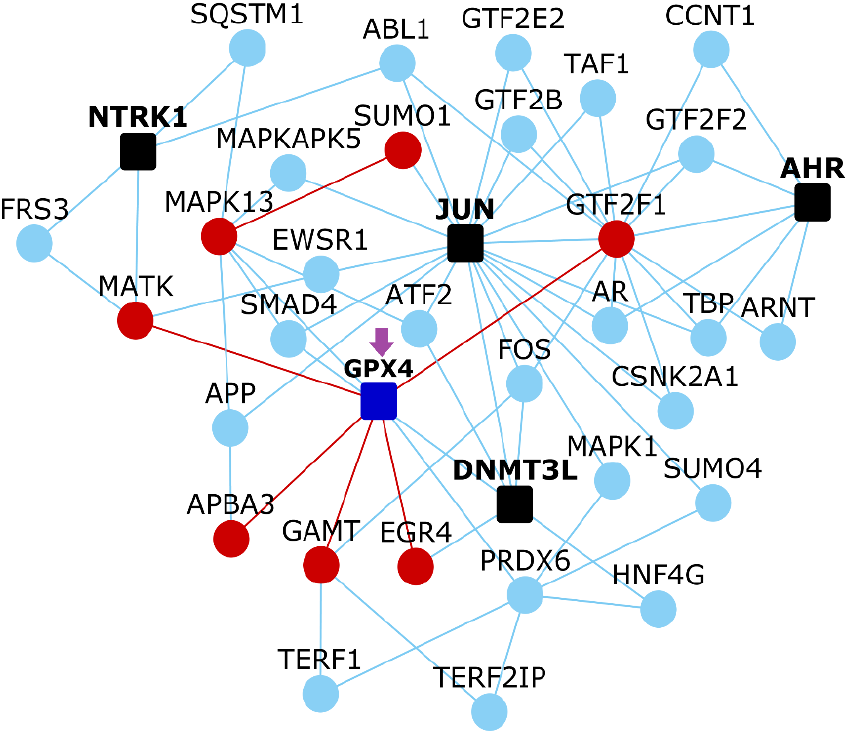
Interactions of GPX4 with genes whose perturbation is known to affect GPX4 expression. This network diagram shows the shortest paths connecting GPX4 with the genes collected from the Knockdown Atlas (black colored nodes), whose perturbation (DNMT3L knockdown/AHR knockout/JUN mutation/NTRK1 overexpression) is known to cause underexpression/overexpression of GPX4. Note that only the PPIs of the 4 perturbed genes that are most closely connected to GPX4 are shown here. Nodes depict proteins and edges depict PPIs. Red and light blue colored nodes denote novel and known interactors respectively. Red and light blue colored edges denote novel and known PPIs respectively. Purple arrows indicate genes associated with heart defects.

*Telomere maintenance/organization* was detected both as global (M1: *Q*-value = 1.16E-04) and fetus-specific (M1: *Q*-value = 1.84E-06) modules. *Positive regulation of neurogenesis* was also detected in global (M7: *Q*-value = 5.88E-03) and fetal (M5: *Q*-value = 5.34E-03) contexts. 6 additional global modules were detected: *cellular response to retinoic acid* (M2: *Q*-value = 1.73E-03), *detoxification* (M3: *Q*-value = 1.73E-03),*peptidyl-threonine phosphorylation* (M4: *Q*-value = 1.73E-03), *regulation of cellular protein localization* (M5: *Q*-value = 3.5E-03), *DNA replication* (M6: *Q*-value =5.35E-03) and *protein autophosphorylation* (M8: *Q*-value = 1.14E-02). 3 additional fetal modules were detected: *negative regulation of myeloid cell differentiation* (M2: *Q*-value = 1.33E-04), *transforming growth factor beta receptor signaling pathway* (M3: *Q*-value = 3.25E-04) and *positive regulation of endopeptidase activity* (M4: *Q*-value = 7.27E-04). 3 novel interactors of GPX4 (EGR4, FUT5 and ZNF197) co-occurred with 2 perturbed genes (NTRK1 and DNMT3L) in the *protein autophosphorylation* global module and in the *neurogenesis* module that was detected in both global and fetus-specific contexts.

### Tissue-elevated gene clusters in the integrated GPX4 landscape

We merged the interactomes described in the previous sections to construct an integrated GPX4 functional landscape containing 342 genes and 461 edges. Then, we sought to isolate clusters of genes showing high expression in specific tissues from this integrated interactome. For this, we generated a data matrix of genes (columns) versus 53 tissues (rows) extracted from GTEx; the cells in the matrix contained log transformed median TPM (transcripts per million) values of gene expression. Principal component analysis (PCA) was used to capture systematic variations underlying this matrix. Using Clustvis,^33^ single value decomposition (SVD) with imputation was applied to this matrix to extract principal components that explain the variance in gene expression observed across the tissues. Unit variance scaling was applied across the matrix. PC1 and PC2 explained 75.1% and 9.6% of the total variance. The log-transformed TPM values were then converted to normalized *Z*-scores. *Z*-scores indicate the number of standard deviations that separate a given TPM value from the mean. This matrix of *Z*-scores was then subjected to hierarchical clustering based on Pearson’s correlation coefficients and the average linkage method.

GPX4 showed high expression in brain tissues and testis. Guo et al. has reported that the human brain and testis exhibit the highest similarity in gene expression patterns among a group of 17 tissues.^34^ This has been attributed to (a) shared biochemical pathways mediated by exocytotic processes and similar receptors between brain and testis tissues, and (b) the involvement of these tissues in human speciation, as a result of which the same set of genes may have been recruited and their expression patterns maintained in both the tissues by evolutionary mechanisms.^35^ We detected a group of 59 genes that clustered with GPX4 and showed high expression in brain tissues and/or testis with/without elevated expression in EBV-transformed lymphocytes and spleen (**Fig. 5**). This cluster included 2 novel interactors of GPX4 that showed high expression in brain tissues, namely, ***EGR4*** and ***MATK***, and ***FUT5*** that showed high expression in testis. Additionally, 5 genes showed elevated expression in both brain and testis tissues (***HHIPL2***, ***MSI1***, COL2A1, MATN1 and MTNR1A (novel interactors shown in bold italics and disease-associated genes shown in bold), 4 genes in only testis (***CHRNB4***, ***SPDYA***, MAGEC1 and RBL1.1), 8 genes in testis, lymphocytes and spleen (**CDKN2A**, ***CLSPN***, ***MSH2***, BLOC1S2, DUSP10, KPNA2, PARP1 and UCHL5) and 39 genes in only brain tissues (***ADAM23***, ***CAMK4***, ***CBS***, ***CDIP1***, ***CHST6***, ***DLG2***, ***DRD2***, ***GOLM1***, ***LY6H***, ***MAPK4***, ***NPTX1***, ***SUSD4***, ARL6IP1, CACNA1A, CALM1, CHAD, COL9A2, COL9A3, DDR1, FEZ1, FGF12, FRS3, GPM6A, HHIP, HID1, HSP90AA1, JPH3, MAG, MAP7, MAPK9, MAPT, MBP, MC4R, PACSIN1, PDIA2, SPP1, TAC1, TERF2IP and YWHAE).

**Figure 5:**
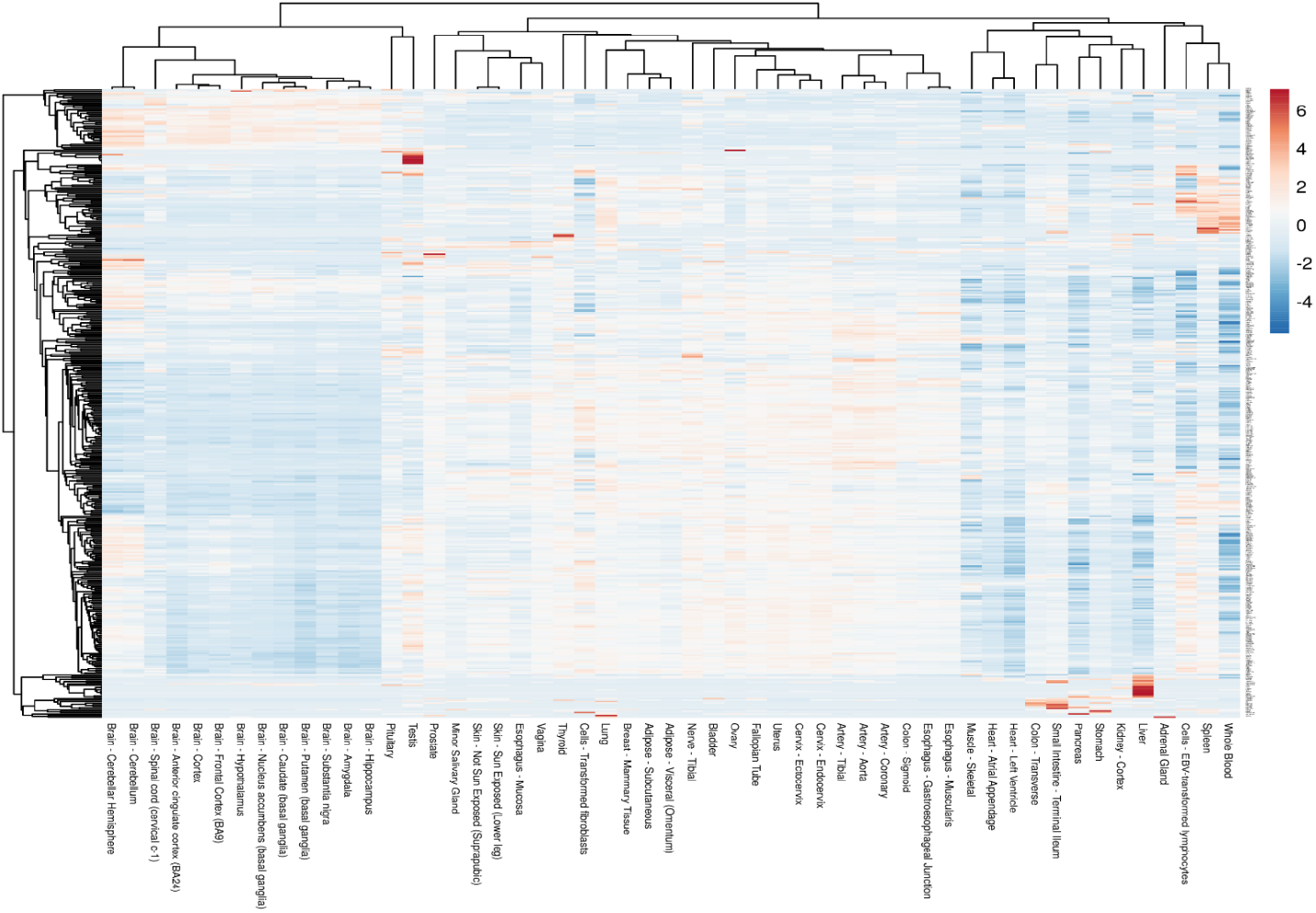
Clustering analysis of genes in the integrated GPX4 interactome. Variations in the expression values of the genes in the integrated GPX4 interactome across 53 tissues have been represented here in the form of a heatmap. The integrated GPX4 interactome contains the shortest paths connecting GPX4 to other genes speculated to be associated with Sedaghatian type spondylometaphyseal dysplasia, genes associated with other spondylometaphyseal dysplasias and phenotypically similar bone dysplasias. Normalized *Z*-scores were computed based on the −log_10_ transformed TPM (transcripts per million) values. *Z*-scores are computed based on the number of standard deviations that separate a given logTPM value from the mean. Clustering was performed using the hierarchical clustering method with average linkage. The dendrograms were derived from the clustering analysis based on computation of Pearson correlation coefficients between the data points. Four gene clusters were detected, namely, a cluster with elevated expression in brain tissues and/or testis with/without elevated expression in EBV-transformed lymphocytes and spleen, another cluster showing elevated expression in whole blood and/or spleen with/without elevated expression in EBV-transformed lymphocytes, and separate liver-elevated and small intestine-elevated clusters.

***GAMT***, a novel interactor of GPX4, showed high expression in the liver, together with a cluster of 15 other genes (***DHODH***, ***SERPINA4***, ***SLC30A10***, ***TM6SF2***, AMBP, APCS, APOB, CRP, HRG, LPA, MST1, NR0B2, ORM2, PLG and TMPRSS6) (**Fig. 5**). 8 genes showed elevated expression in the small intestine (**IHH**, ***CCL20***, ***FCGBP***, HNF4G, MEP1A, MEP1B, POU5F1 and REG3A) (**Fig. 5**). A cluster of 38 genes showed elevated expression in whole blood and/or spleen with/without elevated expression in EBV-transformed lymphocytes (**PSD4**, ***GPR65***, ***IL10RA***, FCGR2B, GCA, ITGA4, ITGB7, LCK, LRG1, LYN, MAP2K6, MX1, NCF2, PDE6G, PTK2B, SRGN, TNF, ***VCAM1***, ***CD3G***, ***RASGRP4***, ***UPP1***, C1QA, C5AR1, CD4, CSFR1, CTSD, CXCR6, DISC1, ELANE, FASLG, GZMM, HCK, HYAL1, ITGA2B, KLRK1, MMP9, NCR1 and TYROBP) (**Fig. 5**).

Enrichment of the brain-elevated genes in axon development, regulation of trans-synaptic signaling, regulation of ion transmembrane transport, protein localization to mitochondrion and neural nucleus development was statistically significant before applying multiple hypothesis correction. The same was true in the case of DNA recombination, aging, regulation of cyclin-dependent protein kinase activity, regulation of DNA metabolic process, mitotic cell cycle phase transition and extrinsic apoptotic signaling pathway for testis-elevated genes, and cell fate commitment, in utero embryonic development, epithelial cell proliferation, mRNA transcription, toxin transport and mesenchymal cell proliferation for small intestine-elevated genes. Enrichment of liver-elevated genes in platelet degranulation and regulation of response to wounding, and lymphocyte/whole blood/spleen-elevated genes in leukocyte differentiation, leukocyte migration, stress-activated protein kinase signaling cascade, adaptive immune response, B cell activation, regulation of inflammatory response and phagocytosis remained statistically significant after multiple hypothesis correction.

We separately inspected the interactome containing the intermediate interactors connecting the 4 genes (AHR, DNMT3L, JUN and NTRK1) whose perturbation affects GPX4 expression, to GPX4, and the known and novel interactors of these 4 genes. We identified a group of 9 testis-elevated genes that clustered with GPX4 (novel interactors shown in bold italics and perturbed genes shown in bold): ***FUT5***, MX1, PDE6G, NCR1, MAGEC1, SUMO4, **NTRK1**, MSX2 and ***FKBP4***.

### Interconnections of GPX4 and genes associated with Joubert syndrome with Jeune asphyxiating thoracic dystrophy

From our Phenogrid analysis, we had noted that Joubert syndrome with Jeune asphyxiating thoracic dystrophy (JATD) showed 64% phenotypic similarity with SMDS. The developmental disorders shared pathophenotypes such as agenesis of corpus callosum, generalized hypotonia, cerebellar hypoplasia and atrial septal defects. Joubert syndrome with JATD shows an amalgamation of several key traits associated with Joubert syndrome such as ataxia-inducing brain stem malformations, hypotonia and cognitive impairment, and skeletal traits characteristic of JATD such as a narrow thorax that leads to respiratory failure, and rib, limb and metaphyseal dysplasia.^36^ We sought to identify whether GPX4 was closely connected to the genes associated with Joubert syndrome with JATD. 3 genes, namely, IFT140, KIAA0586 and CSSP1, which were known to be linked to Joubert syndrome with JATD were extracted from DisGeNET^26^, and the shortest paths connecting these genes (candidates) to GPX4 (target) were identified. IFT140, KIAA0586 and CSSP1 appeared to share 48 intermediate interactors including 14 novel interactors with GPX4 (**Fig. 6a**). The novel interactors included 4 novel interactors of IFT140 (TRAP1, TELO2, IL32 and DNAJA3), 1 novel interactor of KIAA0586 (DACT1), 5 novel interactors of GPX4 (APBA3, GAMT, GTF2F1, MATK and ZNF197) and 4 intermediate interactors (KPNA5, SP100, SNRPC and MAPK13). Additionally, we identified 2 genes from this interactome whose perturbation led to underexpression of Joubert syndrome-associated genes (**Fig. 6a**). It was interesting to note that these genes (TRIM28, ABL1 and BCR1) were known interactors of the novel interactors (ZNF197, GTF2F1 and APBA3) that we had predicted for GPX4. Knockout of TRIM28 led to the underexpression of IFT140, whereas mutations in ABL1 and BCR1 led to the underexpression of KIAA0586.

**Figure 6:**
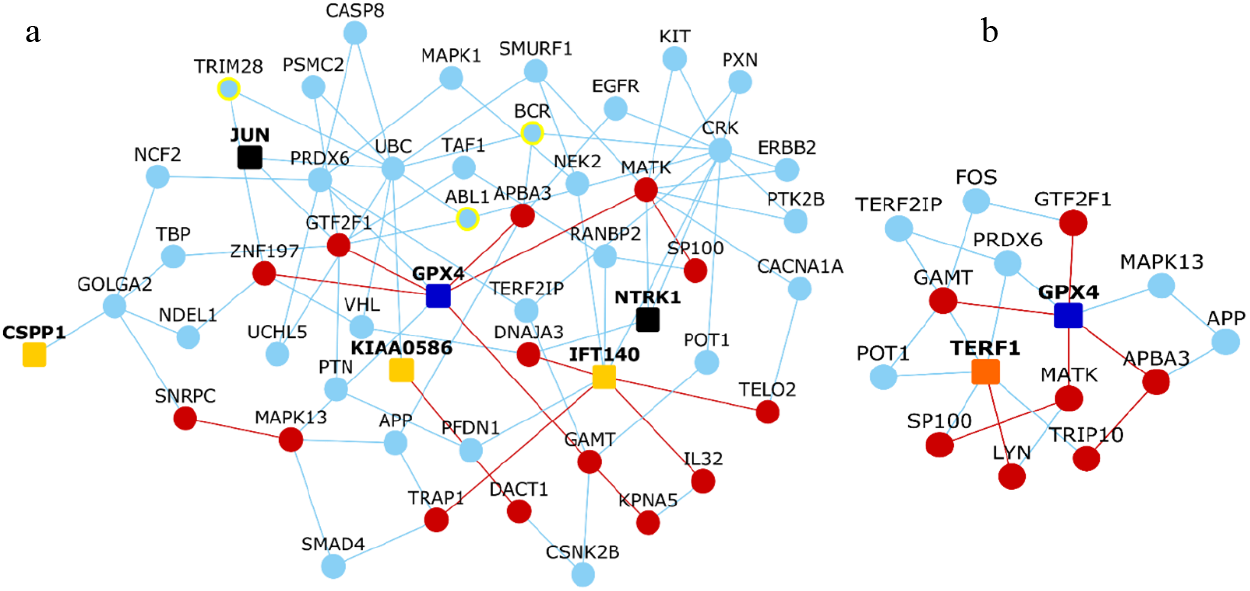
Interactions of GPX4 with ciliary proteins and genes associated with Jourbert syndrome. (a) shows the shortest paths connecting 3 genes associated with Joubert syndrome with Jeune asphyxiating thoracic dystrophy collected from the DisGeNET database (light orange colored nodes) to GPX4; two genes whose perturbation affects GPX4 expression (black colored nodes) and 3 genes (nodes with yellow colored borders) whose perturbation affects 2 Joubert syndrome-associated genes, IFT140 and KIAA0586, can also be found in this network. (b) shows the shortest paths connecting the ciliary protein TERF1 to GPX4. Nodes depict proteins and edges depict PPIs. Red and light blue colored nodes denote novel and known interactors respectively. Red and light blue colored edges denote novel and known PPIs respectively.

Since GPX4 showed close interconnections with the Jourbert syndrome-associated gene IFT140, which is also a ciliary protein, we checked whether GPX4 showed similar interconnections with other ciliary proteins. For this, we examined the shortest paths connecting 165 ciliary proteins^37^ to GPX4. The ciliary protein that showed the closest interconnection with GPX4 was TERF1 (**Fig. 6b**). GAMT, a novel interactor of GPX4, and PRDX6, a known interactor of GPX4, acted as intermediate interactors between GPX4 and TERF1. Overall, GPX4 appeared to be connected to TERF1 through 13 intermediate interactors, including 4 novel interactors of GPX4 (APBA3, GAMT, GTF2F1 and MATK), 1 novel interactor of TERF1 (LYN) and 2 intermediate interactors (TRIP10 and SP100). *Detoxification* was detected both as a global (M4: *Q*-value = 1.26E-03) and fetus-specific (M4: *Q*-value = 1.68E-03) module. Other enriched global modules included *positive regulation of cell development* (M1: *Q*-value = 1.18E-03), *positive regulation of actin filament bundle assembly* (M2: *Q*-value = 1.18E-03), *regulation of G1/S transition of mitotic cell cycle* (M3: Q-value = 1.18E-03), *regulation of JAK-STAT cascade* (M5: *Q*-value = 1.26E-03), *response to growth factor* (M6: *Q*-value = 1.32E-03), *regulation of proteasomal protein catabolic process* (M7: *Q*-value = 1.84E-03) and *establishment of protein localization of organelle* (M8: *Q*-value = 1.94E-03). Other enriched fetus-specific modules included *regulation of binding* (M1: *Q*-value = 4.62E-05), *cellular response to organonitrogen compound* (M2: *Q*-value = 8.91E-04) and *telomere capping* (M3: *Q*-value = 1.05E-03).

### Repurposable drugs for SMDS

We adopted two approaches to identify drugs that may be tested in SMDS from the interactome of SMDS-associated genes compiled from the DisGeNET^26^ database (i.e. GPX4, AGRP, ARNTL, ARTN, LOH19CR1, PSD4 and RPS19). In our first approach, we followed the established methodology of comparing drug-induced versus disease-associated differential expression profiles.^38^ For this, we used a software suite called BaseSpace Correlation Engine (https://www.nextbio.com).^39,40^ This data analysis platform was used because it allows users to study the effect of diseases and/or drugs on thousands of pre-processed publicly available gene expression datasets and has helped to identify drug candidates for diseases such as schizophrenia^41^ (currently undergoing clinical trials^42,43^) and mesothelioma^44^ in the past. We constructed the SMDS drug-protein interactome that showed the drugs that target any protein in the SMDS interactome. 36 drugs targeted 16 proteins in the interactome, including 5 novel interactors (PSMD8, XRCC1, DHODH, GHRL and VHL). We selected 4 gene expression datasets, namely, tibial growth plate hypertrophic zone - Cog mice (chondroplasia) versus wildtype littermates, tibial growth plate hypertrophic zone - Schmid mice (chondroplasia) versus wildtype littermates (GSE30628^45^), skin fibroblasts - Schimke immuno-osseous dysplasia cell line SD60 versus healthy control and skin fibroblasts - Schimke immuno-osseous dysplasia cell line SD8 versus healthy control (GSE35551^46^). Then, we compiled a list of chemical compounds whose differential gene expression profiles (drug versus no drug) were negatively correlated with at least one of the 4 dysplasia-associated differential gene expression datasets (disease versus control). Following this methodology, we identified 7 drugs negatively correlated with chondroplasia and 10 drugs with immune-osseous dysplasia. Although in each case some genes were differentially expressed in the same direction for both the drug and disorder, the overall effect on the entire transcriptome had an anti-correlation. Altogether, we identified 11 drugs as potential candidates that may be tested against SMDS in clinical trials (**Fig. 7a**), namely, anakinra, colchicine, dactinomycin, dexamethasone, fluorouracil, gemcitabine, imatinib, sirolimus, sorafenib, tretinoin and vincristine. Anakinra targets GHRL, a novel intermediate interactor between GPX4 and AGRP. Dexamethasone targets 2 potential SMDS-associated genes, RPS19 and LOH19CR1, and RARA, a known interactor of another SMDS-associated gene called ARNTL. Sorafenib targets a known interactor of GPX4 (MAPK13), a known interactor of ARNTL (HIF1A) and a novel intermediate interactor connecting GPX4 to AGRP (VHL). Both fluorouracil and gemcitabine target SMAD4, a known intermediate interactor connecting GPXE4 to PSD4, and XRCC1, a novel interactor of the SMDS-associated RPS19.

**Figure 7:**
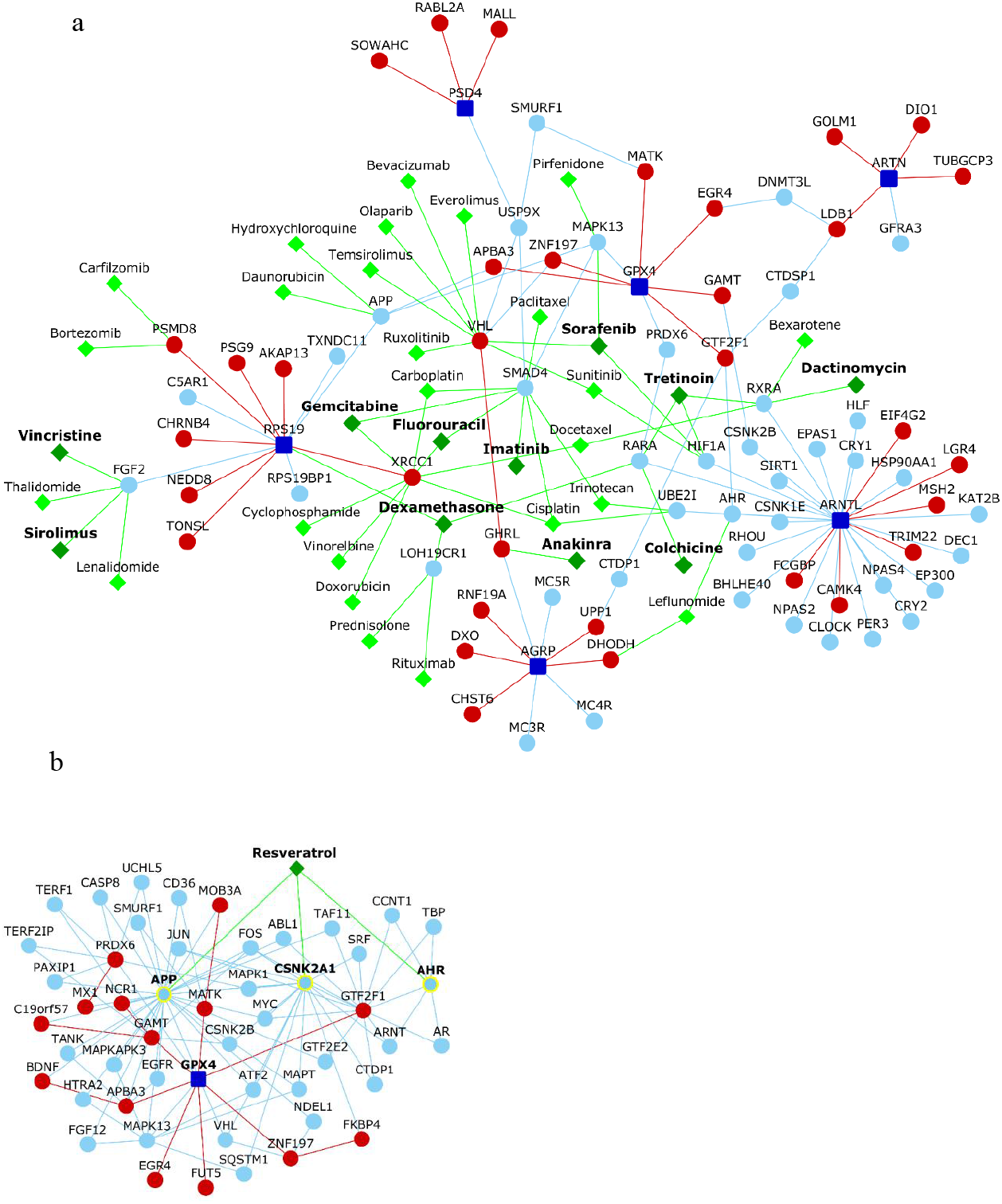
Drug-protein interactome of GPX4. (a) shows the drugs targeting the proteins in the interactome connecting GPX4 to other genes putatively associated with Sedaghatian type spondylometaphyseal dysplasia. Green colored nodes and edges depict drugs and drug-protein interactions respectively. Repurposable drugs identified through comparative transcriptomic analysis have been shown as dark green colored nodes with bold labels. (b) shows the interconnections of GPX4 with 3 targets (nodes with yellow colored borders) of resveratrol. Red and light blue colored nodes denote novel and known interactors respectively. Red and light blue colored edges denote novel and known PPIs respectively.

In our second approach, we compiled a list of drugs that are currently labelled for or are in phase I/II/III clinical trials for different forms of skeletal dysplasias,^47^ and identified their protein targets from Drug Bank.^48^ This yielded a list of 56 proteins targeted by 8 drugs, namely, etidronic acid, risedronic acid, denosumab (osteogenesis imperfecta), resveratrol (pseudoachondroplasia), carbamazepine (Schmid type metaphyseal dysplasia), asfotase alfa (hypophosphatasia), burosumab (X-linked hypophosphatemia) and N-acetylcysteine (diastrophic dysplasia). We examined the shortest paths connecting each of these protein targets to GPX4 and isolated the targets that showed the closest interconnections with GPX4. 3 targets of resveratrol – AHR, CSNK2A1 and APP – appeared to be connected to GPX4 via single intermediate interactors (**Fig. 7b**). Specifically, both AHR and CSNK2A1 were found to be connected to GPX4 via GTF2F1, a novel interactor of GPX4, whereas APP was connected via a known (MAPK13) and a novel (APBA3) interactor of GPX4. We also found that resveratrol induced an expression profile that is negatively correlated with the profile of the Schimke immuno-osseous dysplasia cell line SD60 mentioned before.

## Discussion

SMDS is a severely under characterized skeletal dysplasia driven by GPX4 mutations. In order to expand the functional landscape of this rare and lethal disorder, and expedite the formulation of intervention strategies, we constructed both disease-centric and gene-centric neighborhood networks of GPX4, augmented with novel interactors predicted by the HiPPIP algorithm. Three disease-centric networks were constructed for GPX4, namely, in relation with other putative SMDS-associated genes, SD-associated genes and genes associated with phenotypically similar disorders (**Fig. 1–3**). The GPX4-centric network was constructed to show the interconnections of GPX4 with genes whose perturbation has been known to affect GPX4 expression (**Fig. 4**). Our key findings from these networks were that they were enriched with/contained genes (a) linked to several SMDS pathophenotypes, (b) belonging to tissue-naïve and fetus-specific functional modules, and (c) showing elevated expression in brain and/or testis similar to GPX4. Additionally, we identified 12 drugs that target the neighborhood network of GPX4 and induce gene expression profiles negatively correlated with those associated with chondroplasia and immune-osseous dysplasia.

We used ‘co-membership with GPX4 or its novel interactors in an enriched functional module’ as a criterion to filter genes for experimental dissection of SMDS. Firstly, an integrated network containing the repurposable drugs and the shortest paths of putative SMDS-associated genes, SD-associated genes, genes associated with phenotypically similar disorders and GPX4-affecting genes to GPX4 (collectively referred to as ‘core genes’ henceforth) was constructed. Next, subnetworks containing intermediate PPIs of the genes occurring (along with GPX4 or its novel interactors) in functional modules and the core genes were isolated (**Fig. 8**). Shared phenotypes may reflect interactome-level relationships or similarities in gene function.^49^ In order to facilitate phenotypic-guided investigations of GPX4, we identified 26 genes from these subnetworks that shared at least one phenotype associated with GPX4 (as per data from Mammalian Phenotype Ontology^50^; see **Supplementary File 1**): ADAM23, AGRP, CAMK4, CLSPN, COL2A1, DHODH, DIO1, DNMT3L, DRD2, IHH, IKZF2, KMT2D, LDB1, NTRK1, RASGRP4, RPS19, RXRB, SLC30A10, SPDYA, THRA, TONSL, VHL, XRCC1 and three novel interactors of GPX4 (EGR4, GAMT and ZNF197). LDB1, CAMK4, XRCC1, EGR4 and VHL shared the most number of phenotypes with GPX4.

**Figure 8:**
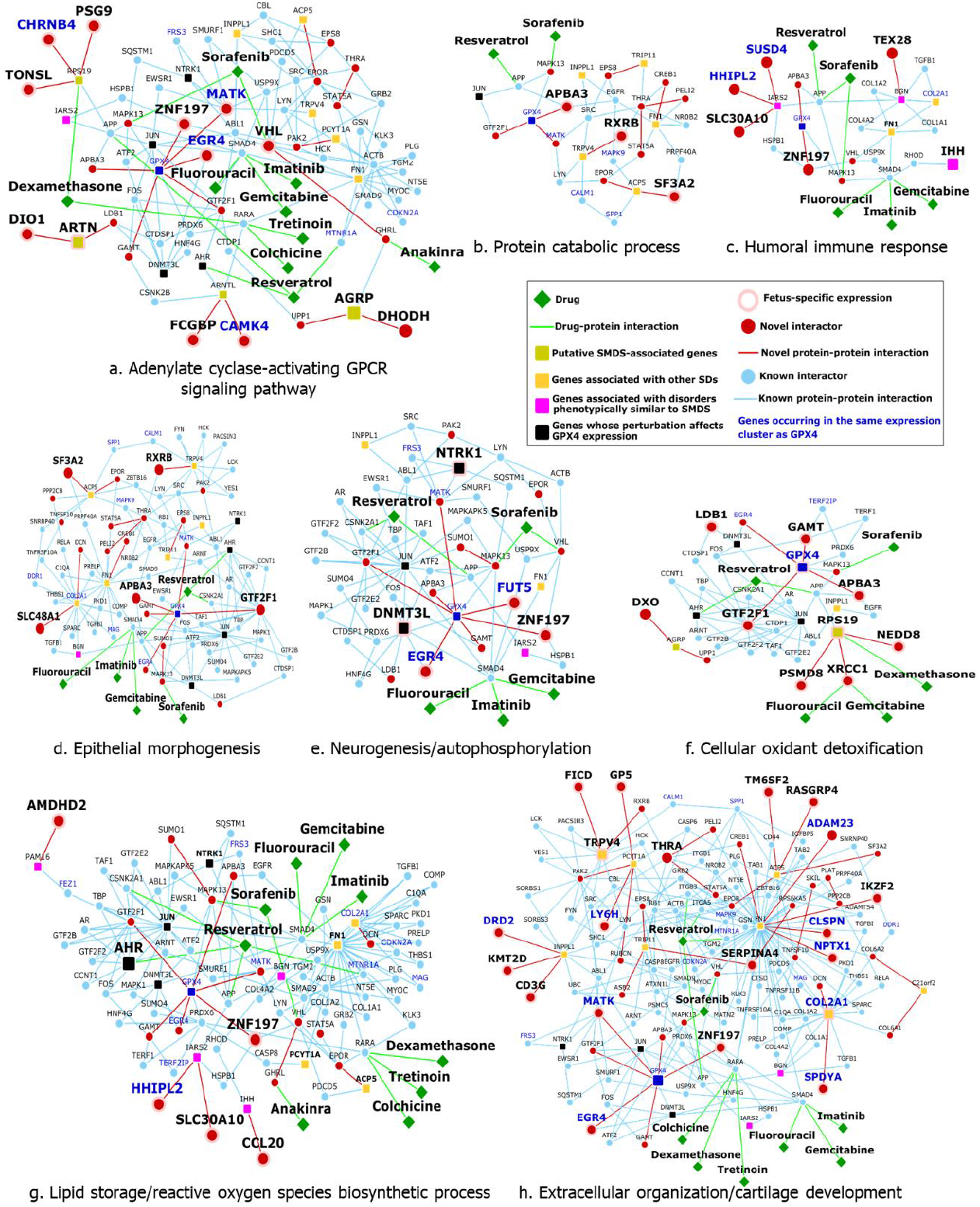
Functional modules with GPX4. The figure shows network diagrams for eight functional modules containing GPX4 or its novel interactors. The network nodes have been annotated as per the legend shown on the right side.

EGR4, which was predicted as a novel interactor of GPX4, is a transcription factor that has been shown to regulate hind brain development in *Xenopus*.^51^ Mice with conditionally deleted XRCC1 (a novel interactor of RPS19) exhibited cerebellar ataxia characterized by reduction in the number of cerebellar neurons and abnormal spike activity in Purkinje cells.^52^ LDB1 is a novel interactor of ARTN (a putative SMDS-associated gene). It interacts with the novel interactors of GPX4 (EGR4 and GTF2F1) via two intermediate interactors (DNMT3L and CTDSP1). LDB1 is an adaptor protein that serves as a critical component of transcription complexes and is involved in the differentiation of various cell types (e.g. hematopoietic cells).^53^ Mutant mice lacking this gene exhibit a range of developmental defects including nterior-posterior patterning, cardiac and foregut defects.^53^ LDB1 has also been shown to influence another gene, LMX1B, mutations in which have been linked to a skeletal dysplasia called the Nail-Patella syndrome.^54^ Accumulation of oxidized lipids leads to a process called ‘ferroptosis’, an iron-dependent/caspase-independent form of apoptosis. Since GPX4 inhibits the accumulation of oxidized lipids by catalyzing their reduction, GPX4 inactivation is critical for ferroptosis. ATF2, a transcription factor that can be found in the *adenylate-cyclase activating GPCR signaling pathway* module (**Fig. 8a**), has been known to inhibit ferroptosis when activated by the JNK1/2 pathway.^55^ Both MAPK13 (a known interactor of GPX4) and CAMK4 (a novel interactor of ARNTL, a putative SMDS-associated gene) have been known to activate ATF2.^56,57^ The exact nature of the functional interaction (e.g. activation/ inhibition) between GPX4 and MAPK13 is unclear. However, if one assumes that GPX4 may inhibit ferroptosis through some indirect action on ATF2, it may be speculated that MAPK13 needs to be activated by GPX4 before it activates ATF2 and influences the inhibition of ferroptosis; note that ATF2 may be independently activated by CAMK4.^58^ Perturbation of GPX4 activity may remove its activational effect on MAPK13, and facilitate ferroptosis. VHL is a known interactor of ZNF197, a novel interactor of GPX4 (**Fig. 8c**). GPX4 has been shown to have an inhibitory effect on elevated VEGFA,^59^ which is generally activated by HIF1A. VHL recruits ZNF197 to inhibit HIF1A transcriptional activity.^60^ Hence, it may be speculated that GPX4 exerts its inhibitory activity on VEGFA, via its direct interaction with the ZNF197-VHL complex. Perturbed GPX4 activity may remove this inhibitory effect on ZNF197-VHL and promote abnormal VEFGA protein expression. Since VEGF signaling is a critical regulator of bone development,^61^ abnormal VEGFA expression may influence the development of skeletal pathophenotypes. Drugs exploiting these potential functional associations of GPX4 on ferroptosis (via MAPK13) and bone development (via ZNF197) could be examined in the context of SMDS. Several other genes in the functional modules have been associated with other dysplasias. Mutations in RPS19 (**Fig. 8f**) and TONSL (**Fig. 8a**) have been associated with hip dysplasia and Sponastrime dysplasia respectively,^62,63^ and KMT2D (**Fig. 8h**) with Kabuki syndrome.^64^ RXRB (**Fig. 8b**) has a regulatory effect on COL2A1, which has been linked to chondrodysplasia, Stickler syndrome and otospondylomegaepiphyseal dysplasia.^65^

We identified 23 genes from the network interconnecting Joubert syndrome-associated genes to GPX4 (**Fig. 6a**), which shared at least one phenotype with GPX4 (**Supplementary File 2**). Most phenotypes were shared by KIT, TRIM28, SNRPC, VHL, CACNA1A, JUN, APP, ABL1 and a novel interactor of GPX4 (GAMT). Additionally, we identified more than 50 genes that showed similar tissue expression patterns as GPX4 and connected GPX4 to other SMDS-associated genes, SD-associated genes and genes whose perturbation affected GPX4 expression. Experimental studies on the mechanistic connections of such genes to GPX4 may provide insights into SMDS etiology.

Comparative transcriptome analysis revealed 11 drugs that induced expression profiles negatively correlated with profiles associated with chondroplasia and immune-osseous dysplasia (**Fig. 7a**). Limited availability of relevant transcriptomic datasets prompted us to use the datasets of these two dysplasias, although they may exhibit etiological distinction from SMDS. We additionally shortlisted resveratrol as a potential drug that may be tested in SMDS due to the proximity of its targets to GPX4 (**Fig. 7b**); resveratrol is currently in phase II trials for pseudoachondroplasia (ClinicalTrials.gov identifier:NCT03866200). The effect of these proposed drugs should be examined in appropriate models.

In summary, our study provides a GPX4-centric network-level view of SMDS, a functional landscape that will allow biologists to prioritize genes, functional modules and drugs for therapeutic interventions in SMDS.

## Methods

### Compilation of gene lists and prediction of novel interactions

SMDS-associated genes, genes associated with X-linked spondyloepimetaphyseal dysplasia, acrocapitofemoral dysplasia, Dyggve-Melchior-Clausen disease, autosomal recessive Megarbane type SMD and genes associated with Joubert syndrome (with JATD) were compiled from the DisGeNET^26^ database. Genes associated with Kozlowski type SMD, spondyloenchondrodysplasia, odontochondrodysplasia, Sutcliffe/corner fractures type SMD, SMD with severe genu valgam/Schmidt type SMD, SMD with cone-rod dystrophy, SMD with retinal degeneration/axial SMD, dysspondyloenchondromatosis, achondrogenesis type 1A, schneckenbecken dysplasia and opsismodysplasia were curated from Warman et al.^31^ Genes whose perturbation (gene knockout/knockdown/mutation/overexpression) affects GPX4 were collected from Knockdown Atlas (BaseSpace Correlation Engine software suite^32^). Novel PPIs of the proteins encoded by these genes were predicted using the HiPPIP model that we developed.^66^ Each protein (say N1) was paired with each of the other human proteins say, (M1, M2,…Mn), and each pair was evaluated with the HiPPIP model.^66^ The predicted interactions of each of the proteins were extracted (namely, the pairs whose score is >0.5, a threshold which through computational evaluations and experimental validations was revealed to indicate interacting partners with high confidence). Previously known PPIs were collected from HPRD (Human Protein Reference Database,^22^ version 9) and BioGRID (Biological General Repository for Interaction Datasets,*^23^* version 3.4.159). The interactome figures were created using Cytoscape.^67^

### Network analysis using LENS

LENS (Lens for Enrichment and Network Studies of human proteins) was used to extract the shortest paths in the human interactome connecting the various sets of genes compiled in this study to GPX4. LENS is a web-based tool that may be used to identify pathways and diseases that are significantly enriched among the genes submitted by users.^33^ The LENS algorithm finds the nearest neighbor of each gene in the interactome and includes the intermediate interactions that connect them. LENS then computes the statistical significance of the overlap of genes in the network and genes with annotations pertaining to pathways, diseases, drugs and GWASs, and reports a *P*-value computed from Fisher’s exact test.

### Identification of functional modules

Functional gene modules were extracted using the HumanBase toolkit^27^ (https://hb.flatironinstitute.org/). HumanBase uses shared k-nearest-neighbors and the Louvain community-finding algorithm to cluster the genes sharing the same network neighborhoods and similar GO biological processes into functional modules. The *P*-values of the terms enriched in the modules are calculated using Fisher’s exact test and Benjamini–Hochberg method.

### Gene expression analysis

Gene expression profiles of 53 postnatal human tissues were extracted from GTEx.^68^ Principal component analysis (PCA) and hierarchical clustering were used to capture relationships among the genes in the various networks constructed in our study (which connect GPX4 to other SMDS-associated genes, SD-associated genes and genes whose perturbation affects GPX4 expression). Log-transformed transcripts per million (TPM) values were assembled into a data matrix containing tissues as rows and genes as columns. PCA was performed with a web-based tool called ClustVis (https://biit.cs.ut.ee/clustvis/).^33^ The data matrix was pre-processed such that 70% missing values were allowed across the rows and columns. The log(TPM) values in the matrix were centered using the unit variance scaling method, in which the values are divided by standard deviation so that each row or column has a variance of one; this ensures that they assume equal importance while finding the components. The method called singular value decomposition (SVD) with imputation was used to extract principal components. In this method, missing values are predicted and iteratively filled using neighbouring values during SVD computation, until the estimates of missing values converge. The data matrix of tissues (rows) and genes (columns) was subjected to hierarchical clustering using the tool called Heatmapper (http://www.heatmapper.ca/)^69^ to identify tissue-based grouping patterns of genes. Pairwise distances in the data matrix were calculated using Pearson correlation and closely linked clusters were identified using the average linkage method. Dendrograms were generated by merging tissues with the smallest distance first, and those with larger distances later. In the average linkage method, the average distance of all possible pairs is considered while clustering.

### Identification of repurposable drugs

The list of chemical compounds whose gene expression profiles correlated negatively with 4 dysplasia expression datasets were compiled using the BaseSpace correlation software (https://www.nextbio.com) (List 1), namely, tibial growth plate hypertrophic zone - Cog mice (chondroplasia) versus wildtype littermates, tibial growth plate hypertrophic zone - Schmid mice (chondroplasia) versus wildtype littermates (GSE30628^45^), skin fibroblasts - Schimke immuno-osseous dysplasia cell line SD60 versus healthy control and skin fibroblasts - Schimke immuno-osseous dysplasia cell line SD8 versus healthy control (GSE35551^46^). Next, we identified drugs that targeted at least one gene in the interactome of SMDS-associated genes (GPX4, AGRP, ARNTL, ARTN, LOH19CR1, PSD4 and RPS19) using Drug Bank (list 2).*^48^* We then compared list 1 and list 2 to identify the drugs that not only target proteins in the interactome but are also negatively correlated with the selected gene expression profiles.

## Author contributions

MKG formulated the project idea to put the interactome-based analysis together motivated by the call “Let’s Find a Cure Together” (https://www.curegpx4.org/). MKG and her students developed PPI prediction and interactome analysis algorithms and tools in prior work. KBK carried out the analysis of the interactomes, presented literature-based evidence, identified repurposable drugs and drafted the manuscript.

## Acknowledgements

There is a call for researchers to come together to find a cure for this rare but debilitating genetic disease (https://www.curegpx4.org/). The little champ Raghav’s smile and the call-to-action through this website motivated us to carry out this analysis. We would be happy to collaborate or provide any additional information in our capacity to researchers who wish to take this work forward, e.g., through experimental validations of the novel PPIs, or testing of the potentially repurposable FDA-approved drugs. MKG’s effort is supported by Department of Biomedical Informatics, School of Medicine, University of Pittsburgh.

## Competing interests

None

## Data and materials availability

Cytoscape files containing the interactomes shown in Figs. 1–4, Fig. 6, Fig. 7, and Fig. 8 have been made available as Supplementary Files 3-6 respectively.

## Supplementary files

**Supplementary File 1:** Functional modules enriched in the networks of putative SMDS-associated genes, SD-associated genes, genes associated with phenotypically similar disorders and GPX4-affecting genes, and the mouse phenotypes shared by genes belonging to these modules.

**Supplementary File 2:** Functional modules enriched in the network of genes associated with Joubert syndrome (with Jeune asphyxiating thoracic dystrophy), and the mouse phenotypes shared by genes belonging to these modules.

**Supplementary File 3:** Cytoscape files containing the networks shown in Figs. 1–4. The networks contain mappings of known (light blue edges) and novel PPIs (red edges) and of known (light blue nodes) and novel interactors (red nodes). The legend for the other nodes are the same as that given in the corresponding figures.

**Supplementary File 4:** Cytoscape files containing the networks shown in Fig. 6. The networks contain mappings of known (light blue edges) and novel PPIs (red edges) and of known (light blue nodes) and novel interactors (red nodes). The legend for the other nodes are the same as that given in the corresponding sub-figures.

**Supplementary File 5:** Cytoscape files containing the networks shown in Fig. 7. Green colored nodes and edges depict drugs and drug-protein interactions respectively. Red and light blue colored nodes denote novel and known interactors respectively. Red and light blue colored edges denote novel and known PPIs respectively.

**Supplementary File 6:** Cytoscape files containing the networks shown in Fig. 8. The legend for the nodes and the edges are the same as that given in the corresponding sub-figures.

## Notes

http://severus.dbmi.pitt.edu/gpx4

### Competing Interest Statement

The authors have declared no competing interest.

### Summary of Updates

Cytoscape files have been added to supplementary files. Author contributions and Acknowledgements edited. MKG affiliation corrected.

